# Alcohol Dependence-Induced Astrocyte Immune Activation in the Nucleus Accumbens

**DOI:** 10.1101/2025.09.12.675849

**Authors:** Joel G. Hashimoto, Regina A. Mangieri, Amanda J. Roberts, Turner Lime, Brett A. Davis, Lucia Carbone, Marisa Roberto, Marina Guizzetti

## Abstract

Astrocytes play many physiological roles in the brain including maintenance of brain homeostasis, modulation of synapse formation and function, and regulation of the blood brain barrier permeability. Upon brain exposure to noxious stimuli, astrocytes can become reactive and activate neuroimmune responses. The nucleus accumbens (NAc) is a critical region involved in reward processing and is integrally involved in the establishment and maintenance of alcohol (ethanol, EtOH) dependence. Here we used the chronic intermittent ethanol – two-bottle choice (CIE-2BC) drinking model to induce EtOH dependence in Aldh1l1-eGFP/Rpl10a mice, which allow the pull-down of astrocyte specific RNA using translating ribosome affinity purification (TRAP) procedure. NAc astrocyte translating RNA and bulk-tissue RNA was analyzed by RNA-Seq to identify genes altered by EtOH dependence or EtOH drinking in astrocytes and the bulk NAc. The number of differentially regulated genes was greater in the astrocyte-specific analysis compared to the bulk-tissue suggesting the cell-type specific approach enables greater resolution of the effects of EtOH. In the astrocyte-specific translatome of EtOH dependent animals, genes related to neuroimmune activation were highly enriched with overall activation of pathways related to interferon and interleukin signaling. In addition, pathways relating to oxidative stress and glutathione related responses were enriched in NAc astrocytes from dependent animals. In contrast, the astrocyte response to EtOH drinking identified pathways related to homeostatic changes. These findings highlight the profound immune activation in NAc astrocytes during EtOH dependence which are distinct from the response to lower levels of EtOH exposure. This study identifies astrocyte pathways and genes involved in the transition from alcohol drinking to alcohol dependence and identify potential novel cell type-specific targets that underlie vulnerability to alcohol use.

## 1. Introduction

The importance of astrocytes in brain function has received increasing levels of attention in recent years. Once thought of as primarily support cells, astrocytes are now known to be integrally involved in synapse formation and function through the production and release of extracellular matrix (ECM) proteins and modulators, and are key components of the neurovascular unit (Kettenmann and Verkhratsky, 2008). Astrocytes respond to CNS damage and disease with changes in gene expression, morphology, and function to become reactive astrocytes. Immune activation is among the most common responses of reactive astrocytes and is an essential component of the brain’s innate immune system (Liss et al., 2025). Different types of brain insult trigger different astrocytic neuroimmune responses and, in the case of chronic diseases, astrocytic responses change with the development of the disease. Alcohol use disorder (AUD) is one of the most prevalent mental health disorders having a lifetime prevalence of nearly 30% in the United States (Grant et al., 2015), and is comorbid with many anxiety-related disorders (Watts et al., 2023). The activation of neuroimmune signaling in AUD has been previously reported (Crews and Vetreno, 2014; Cruz et al., 2023). However, the contribution of astrocytes to AUD-induced neuroimmune responses that increase with the progression of the disorder have been hypothesized but not experimentally explored (Adermark and Bowers, 2016; Guizzetti et al., 2025; Liss et al., 2025).

The understanding of the many functions of astrocytes *in vivo* has been accelerated by new genetic and molecular approaches. One of these is the Translating Ribosome Affinity Purification (TRAP) approach which utilizes a bacterial artificial chromosome (BAC) based transgenic model to express enhanced green fluorescent protein (eGFP) tagged ribosomal protein Rpl10a in specific cell-types using cell-specific promoters (Doyle et al., 2008; Heiman et al., 2008). Immunoprecipitation of cell-type specific eGFP-tagged ribosomes also pulls-down the attached mRNAs that are actively being translated by the ribosomes providing a snapshot of what genes are being translated in a cell-type specific manner. Astrocyte-TRAP mice express an eGFP-tagged Rpl10a protein under the control of the promoter for Aldh1l1, a pan-astrocyte marker (Cahoy et al., 2008), have previously been used in the context of brain development and ageing (Clarke et al., 2018), fetal alcohol spectrum disorders (FASD) (Hashimoto et al., 2025), and the role of connexin 43 in astrocyte immunoreactivity (Boulay et al., 2018), but it has never been applied to study the role of astrocytes in alcohol dependence.

The nucleus accumbens (NAc) is a critical structure for the integration of projections from a variety of brain regions and plays a major role in the regulation of motivated behavior and reward processing (Humphries and Prescott, 2010). The shell of the NAc, together with the central nucleus of the amygdala (CeA), and the lateral portion of the bed nucleus of the stria terminalis (BNST), are the major constituents of the “extended amygdala” (Heimer and Alheid, 1991), a conceptual macrostructure in the basal forebrain that contributes to emotional disturbances, including the initial rewarding effects of alcohol and the negative reinforcement that drives excessive drinking during AUD (Koob, 2008). The NAc is primarily comprised of medium spiny neurons (MSNs), which function as a key node in the convergence of dopaminergic and glutamatergic projections from the ventral tegmental area, prefrontal cortex, hippocampus, and other regions that are involved in the development and maintenance of drug dependence (Britt et al., 2012; Allichon et al., 2021). Alterations in synaptic structure and function in the NAc occur following exposure to substances of abuse, including alcohol/ethanol (ETOH) (Marty and Spigelman, 2012a; Abrahao et al., 2017). In particular, plasticity and metaplasticity (e.g., a loss in the ability to induce long-term depression) of glutamatergic synapses are thought to encode the initiation of EtOH dependence and escalation of voluntary EtOH intake (Jeanes et al., 2011, 2014a; Spiga et al., 2014; Renteria et al., 2016, 2017, 2018; Kircher et al., 2019). In addition, NAc extracellular glutamate tone is increased in dependent animals relative to non-dependent EtOH drinking animals (Griffin III et al., 2014), and contributes to escalation and excessive drinking by dependent animals. Furthermore, NAc structural and functional adaptations can persist for a week or longer into withdrawal (Marty and Spigelman, 2012b; Abrahao et al., 2013; Jeanes et al., 2014b; Peterson et al., 2015; Cannizzaro et al., 2019) and thus likely represent critical mechanisms in not only the development, but also the maintenance, of EtOH dependence.

Despite the abundance of evidence for EtOH-induced NAc adaptations contributing to the escalation of EtOH drinking in EtOH-dependent mice, the role of non-neuronal cells, such as astrocytes, is only beginning to be explored. To understand the role of astrocytes in alcohol dependence, astrocyte-TRAP mice underwent the chronic intermittent EtOH vapor inhalation – two-bottle choice (CIE-2BC) procedure to generate three treatment groups: mice that show increased EtOH intake after chronic EtOH vapor exposure, defined as **EtOH Dependent**; **non-Dependent** control mice with the same voluntary drinking history but without the vapor exposure so they do not escalate their EtOH intake across time; and **EtOH naïve mice** (Becker and Lopez, 2004; Patel et al., 2019; Salem et al., 2024). Astrocyte-TRAP mice exposed to CIE-2BC displayed escalation in EtOH drinking compared to non-Dependent mice which allowed us to compare the astrocyte specific translational changes occurring in EtOH Dependent vs. non-Dependent mice. The inclusion of EtOH naïve mice allowed us to also measure the alterations in astrocyte gene translation caused by EtOH drinking alone, through the comparison of non-Dependent animals to naïve animals. We observe EtOH dependence activates several neuroimmune related pathways in astrocytes. Interestingly, while mice voluntarily drinking EtOH displayed changes in the NAc astrocyte translatome when compared to alcohol-naïve mice, these changes were mostly homeostatic in nature and little neuroimmune activation was observed. These results indicate that the acquisition of a neuroimmune phenotype by astrocytes may be involved in the transition from alcohol drinking to alcohol dependence.

## 2. Methods

### 2.1 Animals

Adult hemizygous B6;FVB-Tg(Aldh1l1-EGFP/Rpl10a)JD130Htz/J mice (Astrocyte-TRAP), generated by Nathaniel Heintz (Doyle et al., 2008; Heiman et al., 2008), were purchased from The Jackson Laboratory (strain # 030247). Animals were acclimated to the animal facility at The Scripps Research Institute for two weeks prior to experimentation. Mice were group housed in individually ventilated caging with Harlan Teklad Aspen Sani-Chip bedding (#7090A) and were fed Envigo Teklad 18% Protein Rodent Diet (#2018). The lights were on a 12/12hr cycle with lights off at 8:00AM. All animal procedures were approved by The Scripps Research Institute Institutional Animal Care and Use Committee following US National Institutes of Health animal welfare guidelines and are in line with the ARRIVE guidelines.

### 2.2 Chronic Intermittent ethanol - Two Bottle Choice (CIE-2BC)

The mouse CIE-2BC model of alcohol dependence-induced escalation of drinking, originally developed by Howard Becker and colleagues (Becker and Lopez, 2004) and adopted by the Roberts lab (Chu et al., 2007; Finn et al., 2007; Warden et al., 2020; Patel et al., 2021; Borgonetti et al., 2023, 2025; Siddiqi et al., 2023; Salem et al., 2024), is widely accepted in the alcohol community and used by at least 20 laboratories worldwide to study the dependence-like alcohol consumption associated with AUD. To acclimate the mice to the EtOH they were singly housed and allowed 24hr access to 15% EtOH in addition to their normal water and food for 4 days (Mon-Fri). After this, mice were returned to their original group housing. Then for an additional 20 days (5 days per week for 4 weeks), 30 min before the lights turn off, mice were singly housed for two hours with access to two drinking tubes, one containing 15% EtOH and the other containing water (i.e. two bottle choice or 2BC). EtOH and water consumption during these 2-hour periods were recorded. Following this baseline period, mice were divided, based on equal EtOH consumptions, into two balanced groups to be exposed to intermittent EtOH vapor (Dependent mice) or control air (non-Dependent mice) in their home cages. The Dependent group was injected with EtOH + pyrazole (alcohol dehydrogenase inhibitor, see below for details) and placed in the chambers to receive intermittent vapor for 4 days (16hr vapor on, 8hr off). Every Thursday, immediately following a 16hr bout of vapor, mice were removed, and tail blood was sampled for blood EtOH (BEC) determination. Target BECs were 175-200 mg/dl. Following the fourth day of exposure, mice were allowed 72 hours of undisturbed time. The mice were then given 5 days of 2-hour access to bottles containing 15% EtOH and water to measure ethanol drinking and preference for the EtOH solution following vapor chamber exposure. The non-Dependent mice were injected with pyrazole in saline at the same time as the vapor groups and received 2BC testing at the same time as the vapor groups. The vapor/air exposure and 5 days of 2 bottle choice testing was repeated another 3 times for a total of 4 rounds of vapor and 2BC testing. Mice were exposed to vapor/air for a final (5^th^) round of CIE and then euthanized within an hour of removal from the chambers for brain collection. Brains were flash frozen in isopentane chilled in a dry ice isopropanol slurry and stored at −80°C until processing. An additional cohort of age-matched EtOH-naïve astrocyte-TRAP animals were left undisturbed in their home cages (i.e. treatment naïve) and were processed as described above for whole-brain dissections.

### 2.3 EtOH Vapor Exposure

EtOH vapor was created by dripping 95% EtOH into a 2000 ml Erlenmeyer vacuum on a warming tray (50°C). EtOH vapor was independently introduced into each sealed chamber (Quad Passive System, La Jolla Alcohol Research, Inc., La Jolla, CA) through a stainless-steel manifold (11 L/min). Mice were kept in their home cages during exposure. Concentrations of EtOH vapor were adjusted by varying the rate at which the EtOH is pumped into the flask and typically range from 22 to 27 mg/liter. To allow finer control over blood alcohol levels the EtOH loading dose and pyrazole dose were varied between 0.875-1.75 g/kg (EtOH) and 34-68.1 g/kg (pyrazole). The dose of pyrazole given to the Dependent mice was also given to the non-Dependent mice. Mice were monitored at least once per day while in the vapor chambers and blood was taken weekly to allow for optimal vapor chamber regulation as well as to best assess each mouse’s health.

### 2.4 Blood EtOH Concentration for Dependent Mice

Approximately 40 µl blood was obtained by cutting 0.5 mm from the tip of each mouse’s tail with a clean surgical blade. Blood was collected in capillary tubes and emptied into Eppendorf tubes containing evaporated heparin and kept on ice. Samples were centrifuged, and plasma decanted into fresh Eppendorf tubes. 5 µl plasma was injected into an Agilent 7820A GC coupled to a 7697A (headspace-flame-ionization). Results were compared with and calibrated using a 6-point serial diluted calibration curve of 300 mg/dl EtOH (Cerilliant E-033).

### 2.5 Translating Ribosome Affinity Purification (TRAP) and RNA isolation

Bilateral brain punch microdissections of the nucleus accumbens (NAc) from frozen whole brains were collected from 300 µm sections using a 1.5 mm brain punch from Bregma 1.7 mm to 0.74 mm (Franklin and Paxinos, 2001). Tissue punches were collected into pre-chilled 1.7 mL microcentrifuge tubes and stored at −80° C until further processing. Frozen brain punches were homogenized in homogenization buffer using a disposable pellet pestle (Kontes) and astrocyte translating RNA was isolated using the TRAP procedure as described (Hashimoto et al., 2025) using anti-GFP antibodies from the Memorial-Sloan Kettering Antibody and Bioresource Core Facility (HtzGFP-19C8 & HtzGFP-19F7) and protein A/G magnetic beads (Thermo Fisher, 88803). RNA was isolated from the input and IP samples using Trizol (Thermo Fisher, Carlsbad, CA) and the Direct-Zol RNA MicroPrep Kit (Zymo Research, Orange, CA). Four batches of TRAP isolations (passes) were carried out, each balanced for treatment group and sex.

### 2.6 RNA-Seq and Analysis

RNA from TRAP and input samples were submitted to the OHSU Integrated Genomics Laboratory (IGL) for sequencing. Libraries were generated by IGL and profiled using the Agilent Tapestation 4200 and sequenced on a NovaSeq 6000 (Illumina). Raw sequencing reads were quality assessed with FastQC v0.11.9 (Andrews, 2010), followed by trimming with Trimmomatic v0.39 (Bolger et al., 2014). Trimmed reads were aligned to the mouse reference genome from Ensembl (GRCm38) with the STAR aligner v2.7.10b (Dobin et al., 2013). The Ensembl annotation gtf file GRCm38.99 was used during alignment to define gene regions and to obtain gene counts from the ReadsPerGene output files from STAR. Unaligned reads were aligned to the EGFP gene using bwa-mem v0.7.17 (Li, 2013). A value representing egfp expression was obtained by taking the number of read pairs mapped to egfp divided by the input read pairs aligned to egfp, multiplied by one million.

For each dataset (RNAseq, TRAPseq, combined), genes with low counts were filtered out prior to downstream analysis with the filterByExpr function from edgeR v3.28.0 (Robinson and Oshlack, 2010). DESeq2 v1.26.0 (Love et al., 2014) was used to perform differential analysis for each dataset. The model used for the contrasts of “Dependent vs non-Dependent”, “Dependent vs naïve”, and “non-Dependent vs naïve” included Sex and TRAP pass as covariates. Additional comparisons were performed separately within each sex, using TRAP pass as a covariate. Significant regulation was determined by multiple-comparison adjusted p-values < 0.05. Finally, comparisons for “IP vs input” were performed on the combined dataset within each treatment (Dependent, non-Dependent, naïve) using sex and TRAP pass as covariates. For IP vs Input comparisons, significant enrichment in the IP (astrocyte) fraction was determined by an adjusted p-value < 0.05 and a log_2_ fold-change > 1.

### 2.7 TRAP q-RTPCR

Confirmation of cell-type specific enrichment by the TRAP procedure and TRAP RNA-Seq results was carried out using qRT-PCR on a subset of input and IP samples used for the RNA-Seq as previously described (Hashimoto et al., 2025; Kawa et al., 2025). Diluted aliquots of input and IP RNA were analyzed using the Luna Universal One-Step RT-qPCR Kit (NEB) on the CFX96 (Bio-Rad) thermocycler with normalization to total RNA using the Quant-it RiboGreen kit (ThermoFisher). Primer sequences for *Il33*, *Aldh1l1*, *Tubb3, Itgam*, and *Mbp* were previously described (Talabot-Ayer et al., 2012; Hashimoto et al., 2025).

### 2.8 Bioinformatic Analysis

Gene category enrichment analysis was carried out using Ingenuity Pathway Analysis (IPA, Qiagen) to determine enriched canonical pathways. Canonical pathway enrichment analysis produces p-values corresponding to the significance of the enrichment, enrichment ratio corresponding to the number of enriched genes in the analyzed dataset divided by the total number of genes in the pathway, and activation z-score which is a prediction of pathway activation or inhibition based on the direction of regulation of genes in the dataset.

### 2.9 Statistical Analysis

For the BEC data (obtained only from the Dependent animals), effects of sex, CIE week (i.e. the 5 different weeks of CIE), and sex by CIE week interaction were assessed by two-way repeated measure ANOVA. Parameters measured during the 2BC drinking (EtOH g/kg, body weight, EtOH mL, and water mL) were analyzed using a linear mixed effects model to account for the multiple measurements within each week of the 2BC drinking in the R package lme4 (Bates et al., 2015) using intercept-only models as previously described (Goeke et al., 2018). Effects of sex, EtOH vapor exposure (CIE), and sex by EtOH vapor exposure interactions were assessed by likelihood ratio tests with p-values generated by chi-squared test, where significance is determined as p < 0.05.

## 3 Results

### 3.1 CIE-2BC induced EtOH dependence results in escalation of drinking

Astrocyte-TRAP mice underwent the CIE-2BC procedure which involves cycles of 2BC drinking for 2 h/day for 5 days and chronic EtOH vapor exposure for 16 h/day for 4 days (Dependent mice; Figure 1A). Non-Dependent animals undergo the same procedure but are not exposed to EtOH vapor (see Methods). Following each vapor exposure, blood ethanol concentrations (BECs) were determined (Figure 1B). Analysis of BECs show no effects of sex or sex by CIE week interaction but a significant effect of CIE week (F_(4)_ = 10.2, p = 8.5×10^−7^). The hallmark for 2BC-CIE induced EtOH dependence is escalation of drinking in EtOH vapor exposed animals (Dependent) compared to non-Dependent animals. After the 4^th^ CIE (CIE4, Figure 1C), EtOH vapor exposed animals showed increased EtOH drinking (g/kg) compared to controls (χ^2^_(1)_ = 22.4, p = 2.2×10^−6^) as well as sex differences in EtOH drinking (χ^2^_(1)_ = 249.3, p = 3.7×10^−56^). The drinking level for each animal after the 4^th^ CIE are presented in Figure 1D to show the individual variation in drinking levels with female Dependent animals drinking towards the high end and male controls showing lower EtOH drinking.

**Figure 1.**
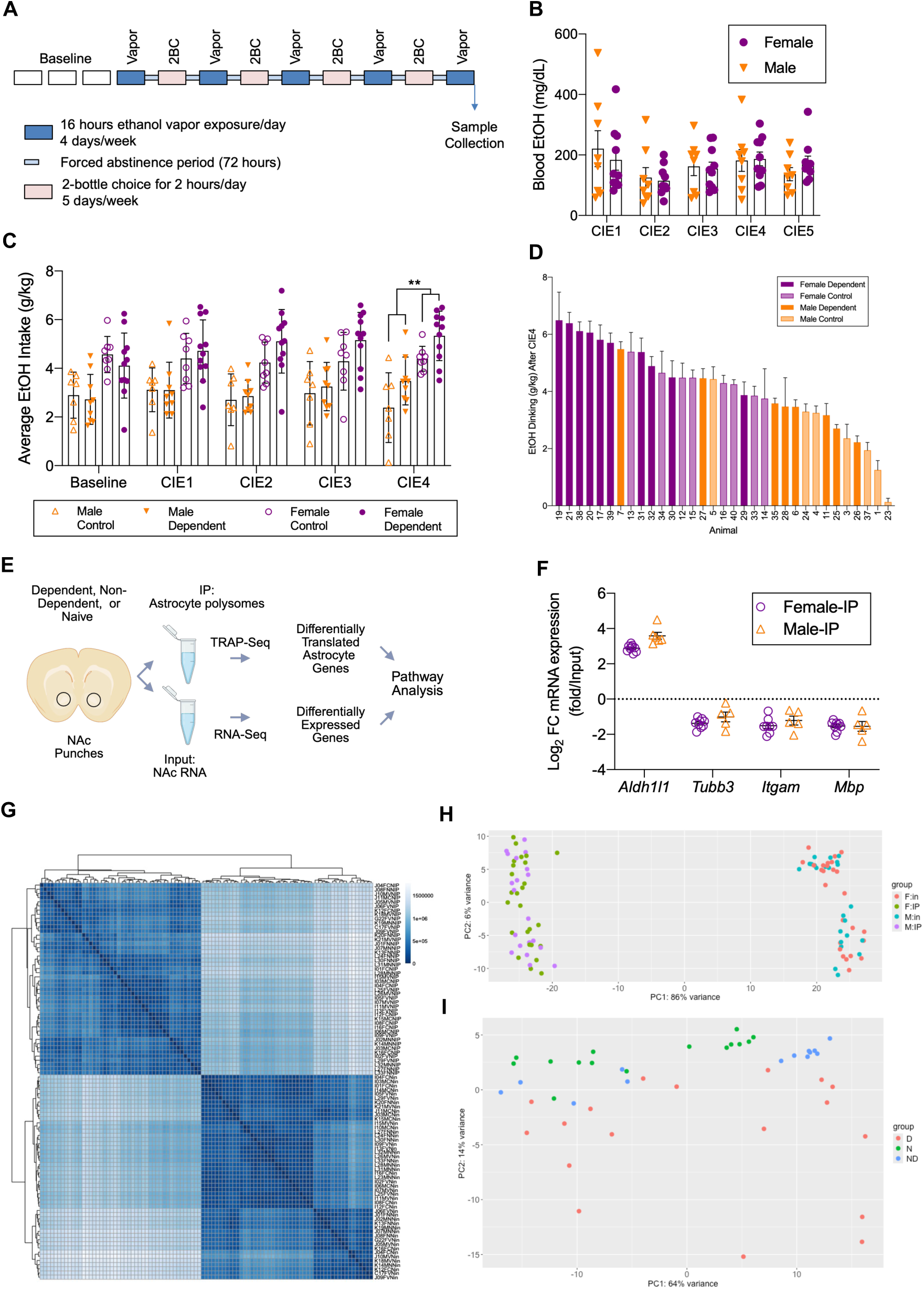
**A.** Schematic of CIE-2BC procedure to induce ethanol dependence. **B**. Blood ethanol concentrations following each CIE vapor exposure. **C.** Average drinking during baseline and after the first four rounds of CIE (2BC drinking is not done after the 5th CIE exposure). **D.** Individual animal drinking after the 4^th^ CIE exposure showing individual variability of drinking levels. **E.** Schematic of sample and data processing for TRAP-Seq and RNA-Seq. Brain punches from Dependent, non-Dependent, and naïve animals were processed using the TRAP protocol. RNA from both fractions were sequenced to identify astrocyte differential translation and bulk tissue differential expression between relevant treatment groups. In addition, IP (immunoprecipitated astrocyte RNA) and input (bulk-tissue) samples were compared to identify genes enriched in astrocytes. **F.** A subset of IP and input samples were analyzed by qRT-PCR with astrocytic (*Aldh1l1*), neuronal (*Tubb3*), microglial (*Itgam*), and oligodendrocyte (*Mbp*) cell-type specific primers. *Aldh1l1* shows robust enrichment in IP compared to input while the other marks show de-enrichment in IP samples compared to inputs. Treatment condition was collapsed in this graph for easier interpretation. **G.** Hierarchical clustering of of NAc RNA-Seq samples from Dependent, non-Dependent (control), and treatment naïve animals from both input (bulk-tissue) and immunoprecipitated (IP) astrocyte translating RNA. The main separation is between IP and input samples with the darker blues indicating IP samples clustering together in the top left portion of the plot and the input samples clustering together in the lower right portion of the plot. Sample names are on the right side of the cluster with the first 3 characters the animal, the 4^th^ character the sex, the 5^th^ character the treatment (V: Dependent, C: non-Dependent, N: naïve), the 6^th^ character the tissue (N: NAc), and the 7^th^-8^th^ character the fraction (IP, in: input) so the first sample, J04FCNIP, is animal J04, a female control NAc IP sample. **H.** PCA plot of all samples showing distinct separation of IP and input samples. Samples are colored based on sex and fraction, with female input (F:in) samples red, female IP (F:IP) samples green, male input (M:in) samples blue, and male IP (M:IP) samples purple. **I**. PCA of only the IP samples only shows separation of Dependent (D) samples from, non-Dependent (ND) and naïve (N) samples primarily along PC2.

### 3.2 EtOH alters NAc transcriptome and astrocyte translatome

To understand how NAc astrocytes respond during EtOH dependence, we utilized the astrocyte-TRAP transgenic mouse model (expressing an eGFP-tagged ribosomal protein, Rpl10a, under the control of the promoter for the astrocyte marker *Aldh1l1*) which enables the isolation of astrocyte enriched translating RNA from brain tissues (Doyle et al., 2008; Heiman et al., 2008). Translating astrocyte mRNA (after immunoprecipitation, IP, with an eGFP antibody) and mRNA from an aliquot of the input (prior to IP) were isolated and sequenced (Figure 1E). Enrichment of astrocyte marker *Aldh1l1* and depletion of non-astrocytic markers *Tubb3, Itgam,* and *Mbp* in IP (astrocyte) versus Input samples are shown in Figure 1F, similar to what we previously reported in the developing hippocampus (Goeke et al., 2022; Hashimoto et al., 2025). Input and astrocyte translating RNA from Dependent, non-Dependent, and naïve NAc samples were sequenced by the OHSU IGL core facility. The average RNA integrity number (RIN) was 7.6; RINs from individual samples analyzed by RNA-seq are shown in Supplemental Table 1.

Hierarchical clustering of samples shows clear separation of input versus IP samples (Figure 1G). Similarly, principal component analysis (PCA) separates the IP vs input samples along principal component (PC) 1 (86% of variance, Figure 1H). PCA on the IP samples alone shows some separation based on treatment (Figure 1I) with the Dependent group showing the most overall variability. In addition, we analyzed the expression of eGFP to determine if the treatments or sex of the animals impacted the expression of the transgene and found no effects of sex, treatment or interaction of treatment by sex in the IP or input fractions (data not shown).

The number of differentially translated (DT; astrocyte or IP fraction) and differentially expressed (DE; input or bulk-tissue fraction) genes based on the three comparisons (Dependent vs. naïve, Dependent vs. non-Dependent, and non-Dependent vs. naïve) in each of the two fractions (IP and input) are summarized in Table 1. In addition, genes that have higher expression (enriched) in astrocytes compared to the bulk tissue expression, suggesting of primarily astrocytic functions, were identified using an adjusted p-value < 0.05 and a log_2_ fold-change > 1 between the IP and input fractions in any of the groups (Dependent, non-Dependent, naïve) are reported in Supplemental Table 2, as we have done previously (Hashimoto et al., 2025). The number of differentially translated or expressed genes across treatments that are enriched in astrocytes are summarized in the final column of Table 1. The full list of DT and DE genes are available in Supplemental Table 3. When looking at the overall effects of EtOH with sexes combined, the most DT and DE genes were identified in the Dependent vs naïve comparison. This is not surprising as these are the most different treatments, i.e. animals with no alcohol exposure (naïve) vs. animals that have undergone voluntarily alcohol drinking and have also been exposed to ethanol vapor (Dependent).

**Table 1.**
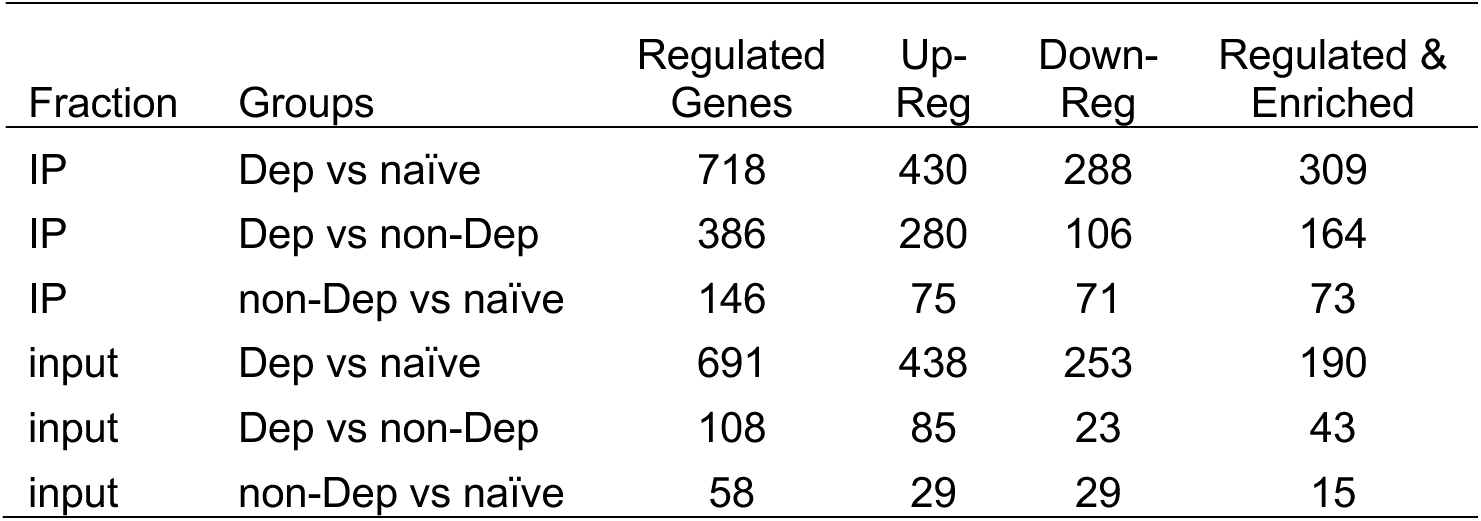
Number of Differentially Translated and Differentially Expressed Genes. Fraction refers to the astrocyte (IP) translating RNA-Seq or bulk-tissue (input). Group refers to the comparison being made (Dep: Dependent; Non-Dep: non-Dependent; naïve). Regulated genes were determined with an adjusted p-value < 0.05. Up-Reg (or Down-Reg) refers to the number of genes up-regulated (or down-regulated) in the first named group in each comparison (i.e. Dep vs naïve, Up-Reg genes are higher in Dependent animals compared to naïve animals). Regulated & Enriched refers to the number of regulated genes that were also significantly higher (>1 log_2_ fold-change) in the astrocyte (IP) fraction compared to the bulk-tissue (input).

| Fraction | Groups | Regulated Genes | Up-Reg | Down-Reg | Regulated & Enriched |
| --- | --- | --- | --- | --- | --- |
| IP | Dep vs naïve | 718 | 430 | 288 | 309 |
| IP | Dep vs non-Dep | 386 | 280 | 106 | 164 |
| IP | non-Dep vs naïve | 146 | 75 | 71 | 73 |
| input | Dep vs naïve | 691 | 438 | 253 | 190 |
| input | Dep vs non-Dep | 108 | 85 | 23 | 43 |
| input | non-Dep vs naïve | 58 | 29 | 29 | 15 |

Of particular significance are the changes in translation (IP) and expression (input) (386 and 108 genes, respectively) observed in samples from Dependent vs non-Dependent mice, as these are likely to be relevant to the understanding of the biology behind the transition from recreational drinking to AUD. The comparison between non-Dependent vs naïve samples elicited the fewest changes, both in the IP and input. In all comparisons, we observed more changes induced by 2BC or CIE-2BC (EtOH drinking or EtOH vapor exposure plus EtOH drinking) in the astrocytes (IP) DT genes than in the input DE genes, which we have observed previously under different conditions (Hashimoto et al., 2025) and is consistent with the idea that cell-type specific approaches allow for increased mechanistic insights.

The number of astrocyte DT genes that overlap based on the different comparisons are 147 in Dependent vs. naïve and Dependent vs Non-Dependent, 83 in Dependent vs. naïve and non-Dependent vs. naïve, 16 in Dependent vs. non-Dependent and non-Dependent vs. naïve, and 1 gene (*Gstm1*), a gene enriched in astrocytes and encoding for glutathione S-transferase mu-1, was up-regulated by EtOH in all 3 comparisons (Figure 2A). The astrocytic dysregulation of *Gstm1* by EtOH is of note as this gene has been shown to increase inflammatory signaling in astrocytes resulting in astrocyte mediated microglia activation during brain inflammatory responses (Kano et al., 2019). As member of the glutathione S-transferase family of enzymes, *Gstm1* is also involved in the detoxifying and antioxidant effects of glutathione as also discussed below (Aloke et al., 2024).

**Figure 2.**
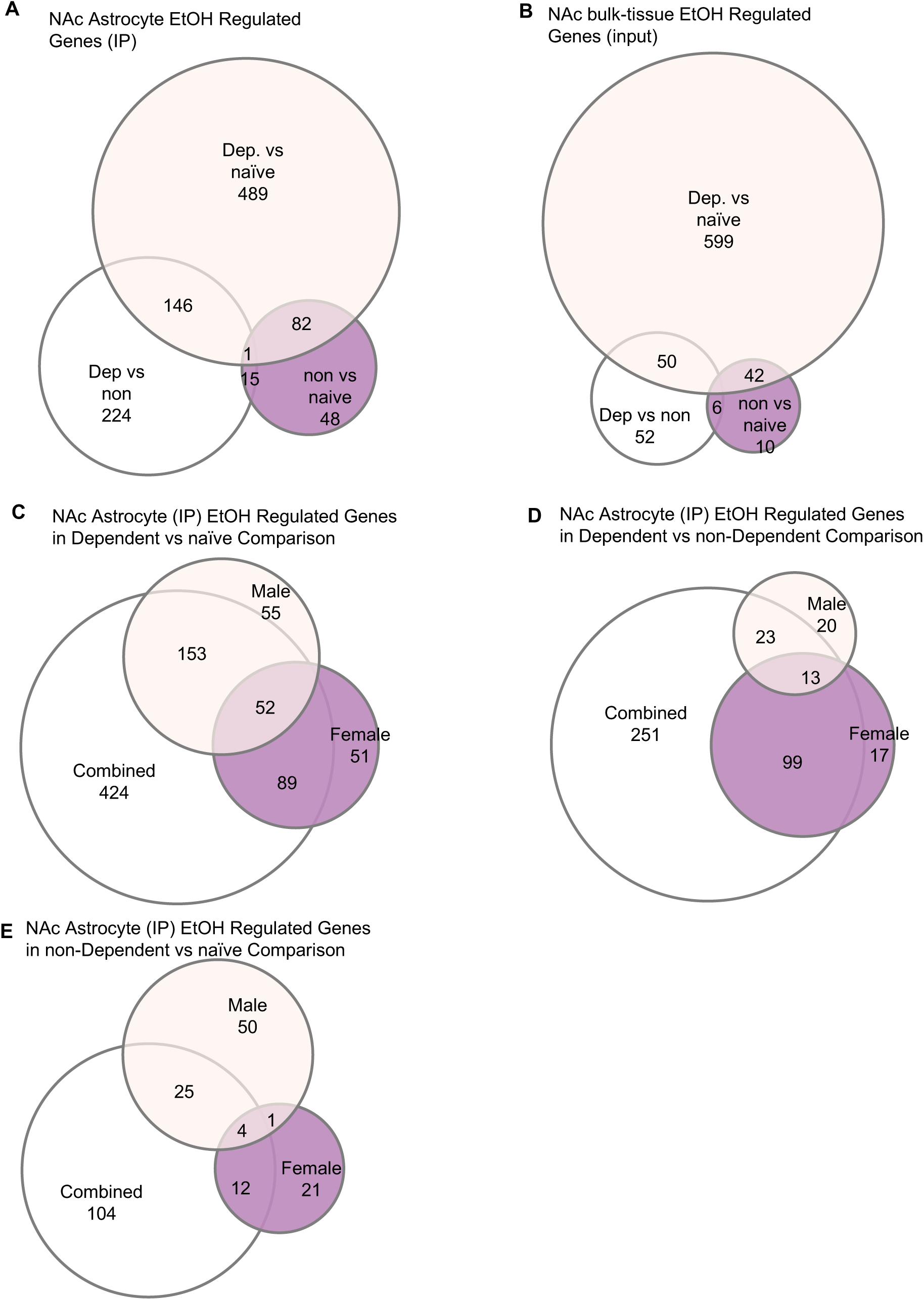
Venn diagrams showing overlap in different comparisons. **A.** Overlap of significantly regulated genes in the Dependent (Dep.) vs naïve, Dependent vs non-Dependent (non), and non-Dependent vs naïve comparisons in the IP (astrocyte) samples. **B.** Overlap of significantly regulated genes in the Dependent vs naïve, Dependent vs non-Dependent, and non-Dependent vs naïve comparisons in the input (bulk-tissue) samples. **C.** Overlap of significantly regulated genes in the Dependent vs naïve comparison when looking at sexes combined, females only, and males only in the IP fraction. **D.** Overlap of significantly regulated genes in the Dependent vs non-Dependent comparison when looking at sexes combined, females only, and males only in the IP fraction. **E.** Overlap of significantly regulated genes in the non-Dependent vs naïve comparison when looking at sexes combined, females only, and males only in the IP fraction.

The number of NAc DE genes that overlap in the comparisons: Dependent vs. naïve and Dependent vs. non-Dependent is 50; Dependent vs. naïve and non-Dependent vs. naïve is 42; Dependent vs. non-Dependent and non-Dependent vs. naïve is 6. There were no genes that were regulated in all comparisons in the input (bulk-tissue) fraction (Figure 2B).

The overlap of astrocyte genes regulated when including both males and females versus females alone and males alone are summarized in Figure 2C-E. There is a large proportion of overlapping genes in the Dependent vs naïve and Dependent vs non-Dependent comparisons with less overlap in the non-Dependent vs naïve comparison. This may be due to the more modest effects of 2BC drinking (non-Dependent) compared to CIE-2BC (Dependent), the inter-animal variability in 2BC drinking (Figure 1D), or the timing of when the tissue was collected (after the 5^th^ vapor exposure when the non-Dependent animals had no EtOH for over a week).

### 3.3 EtOH dependence alters astrocyte neuroimmune signaling

Ingenuity Pathway Analysis (IPA) was used to determine pathway overrepresentation in genes that were regulated by EtOH and to identify pathway activation or deactivation based on known gene relationships and direction of regulation (Figure 3A & B). The following discussion will focus on genes that are regulated in astrocyte samples (IP) in the Dependent vs non-Dependent comparison as it allows us to distinguish between genes regulated by voluntary EtOH drinking (2BC) and genes that are regulated during the process of establishing EtOH dependence (CIE-2BC).

**Figure 3.**
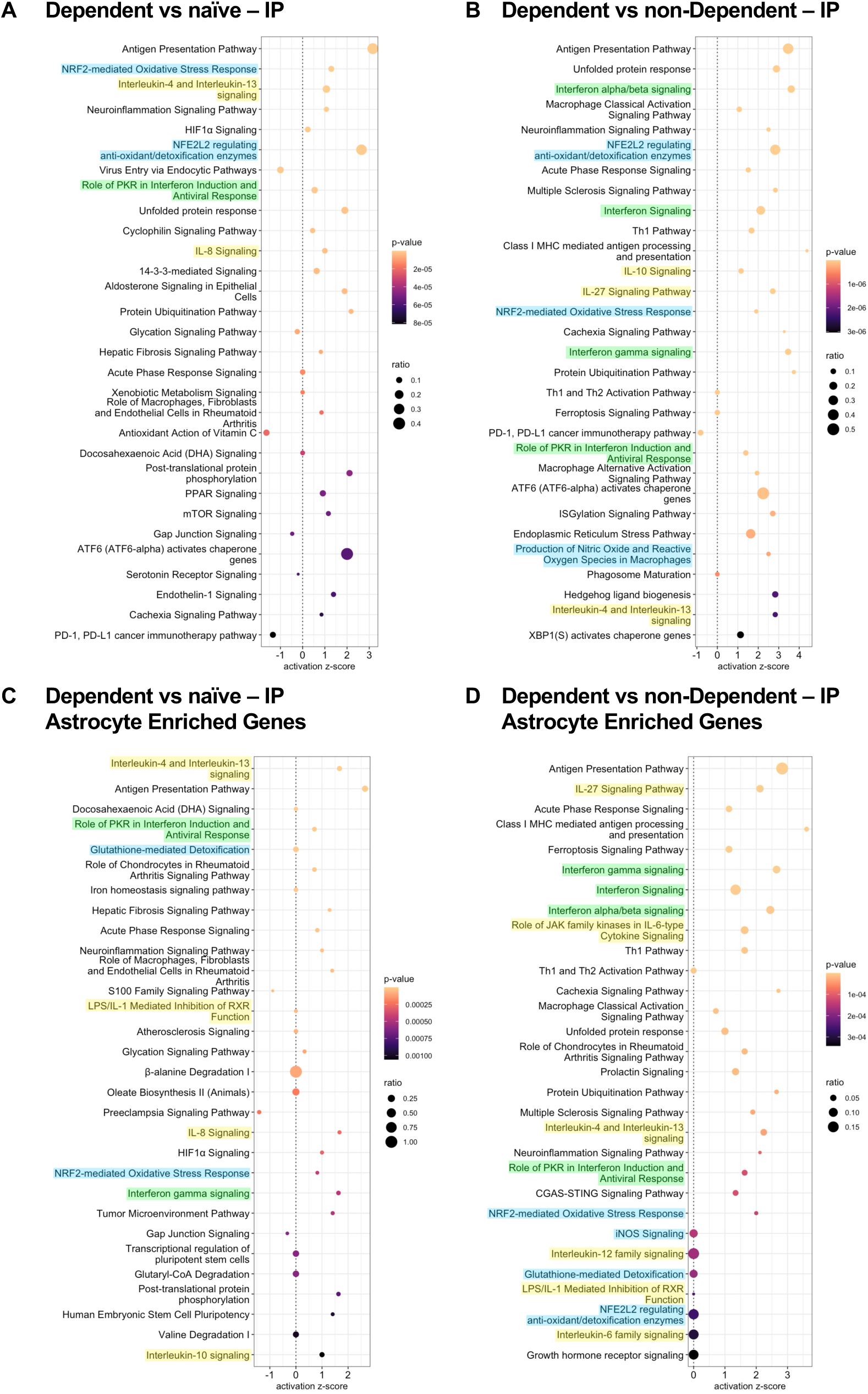
Top 30 enriched canonical pathways from IPA of Dependent vs naïve (**A**) and Dependent vs non-Dependent comparisons (**B**) in the IP (astrocyte) samples. Significantly differentially translated genes in the astrocyte fraction that were also enriched in astrocytes were analyzed for pathway overrepresentation in the Dependent vs naïve comparison (**C**) and the Dependent vs non-Dependent comparison (**D**). Pathways highlighted in yellow relate to interleukin signaling, green highlights relate to interferon, and blue highlights relate to oxidative stress and glutathione pathways. In each panel, the x-axis shows the activation z-score which assigns positive or negative activation based on known pathway dynamics and the regulation of genes enriched in the pathway.

A first important observation is that among the top 30 canonical pathways based on enrichment p-value in the Dependent vs non-Dependent IP comparison there are numerous pathways that relate to immune functions and that most of these pathways have a positive z-score indicating activation of these neuroimmune pathways in astrocytes during the transition to alcohol dependence (Figure 3B). The complete list of IPA canonical pathways significantly overrepresented (p-value < 0.01) in the Dependent vs non-Dependent IP is reported in Supplemental Table 4.

One neuroimmune function activated in Dependent animals involves interferon signaling, as indicated by the fact that among the top 30 significant pathways, “Interferon alpha/beta signaling”, “Interferon Signaling”, and “Interferon gamma signaling” were listed (Figure 3B; interferon-related pathways are highlighted in green). In the complete list of significantly overrepresented IPA canonical pathways (Supplemental Table 4) there are 6 pathways relating to interferon; these pathways and the regulated genes within them are summarized in Table 2 and Figure 4A.

**Figure 4.**
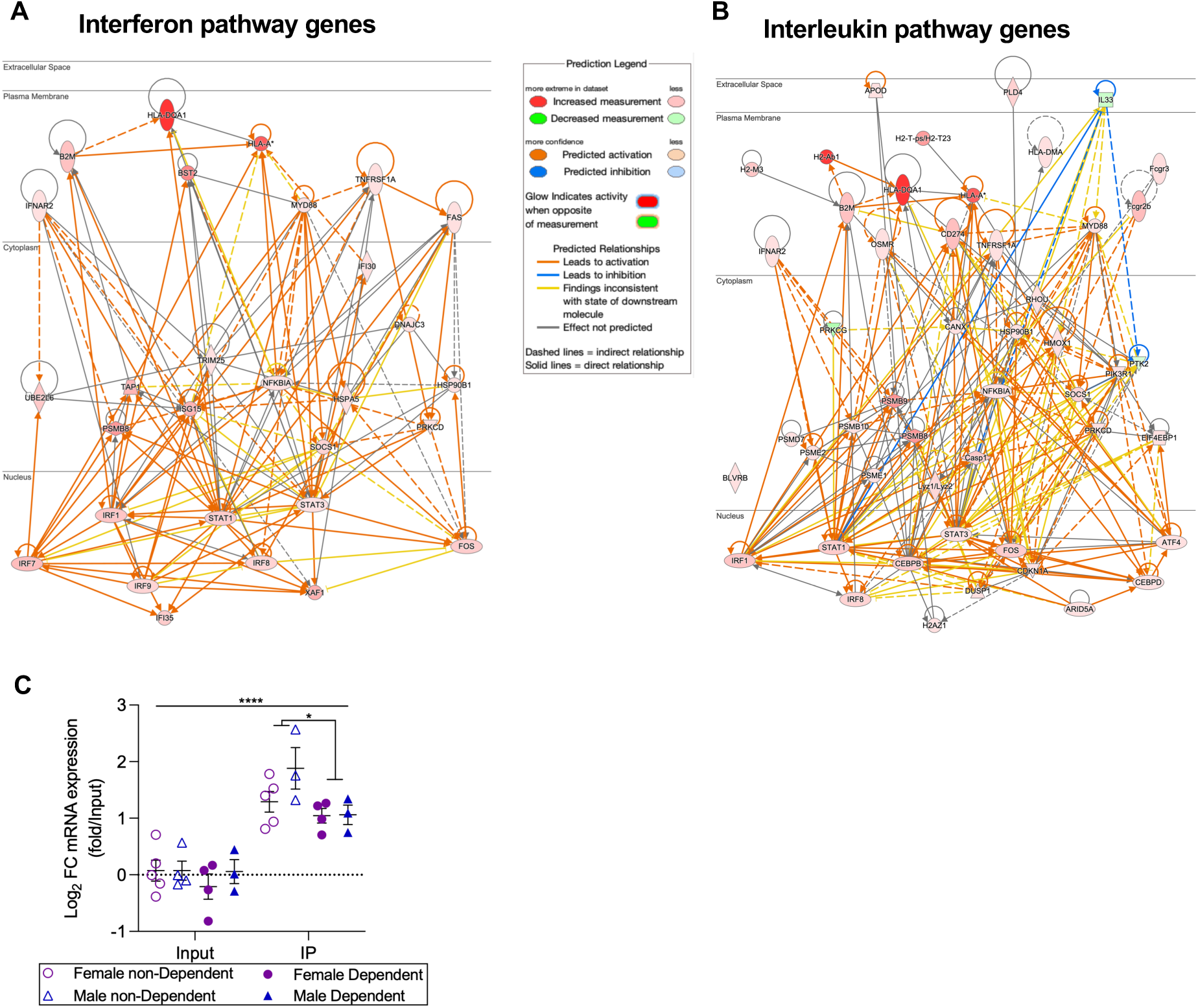
Interferon and Interleukin related genes in NAc Astrocytes from EtOH Dependent mice. **A.** Interferon related pathways were significantly overrepresented in Dependent vs non-Dependent NAc astrocytes. Genes and pathways summarized in Table 2 were used to create a network of known direct and indirect interactions using IPA. **B.** Interleukin related pathways were significantly overrepresented in Dependent vs non-Dependent NAc astrocytes. Genes and pathways summarized in Table 3 were used to create a network in IPA. **C.** Confirmation by qPCR of RNA-seq data on *Il33* expression and translation. *Il33* down-regulation in the IP fraction from the NAc of Dependent vs non-Dependent mice (*, p = 0.038); *Il33* enrichment in astrocytes (IP fraction) compared to input (bulk tissue; ****, p < 0.0001).

**Table 2.** Interferon Related Pathway Genes. Gene symbols in bold indicate the gene is enriched in the astrocyte (IP) fraction. Categories column refers to the following IPA canonical pathways: 1) Interferon alpha/beta signaling; 2) DDX58/IFIH1-mediated induction of interferon-alpha/beta; 3) Interferon gamma signaling; 4) Interferon Signaling; 5) Interferon Signaling; 6) Role of PKR in Interferon Induction and Antiviral Response.

| Symbol | Gene Name | Log<sub>2</sub> FC | adj. p-value | Categories |
| --- | --- | --- | --- | --- |
| **Dnajc3** | DnaJ heat shock protein family (Hsp40) member C3 | 0.393 | 8.31E-04 | 6 |
| Ifnar2 | interferon (alpha and beta) receptor 2 | 0.409 | 2.30E-03 | 1, 5, 6, 4 |
| **Stat3** | signal transducer and activator of transcription 3 | 0.428 | 1.97E-04 | 5, 6 |
| **Tnfrsf1a** | tumor necrosis factor receptor superfamily, member 1a | 0.443 | 7.63E-05 | 6 |
| **Myd88** | myeloid differentiation primary response gene 88 | 0.452 | 2.68E-02 | 6 |
| Nfkbia | nuclear factor of kappa light polypeptide gene enhancer in B cells inhibitor, alpha | 0.496 | 1.56E-02 | 2, 6 |
| Hsp90b1 | heat shock protein 90, beta (Grp94), member 1 | 0.499 | 1.29E-03 | 6 |
| Hspa5 | heat shock protein 5 | 0.621 | 1.40E-05 | 6 |
| **Trim25** | tripartite motif-containing 25 | 0.637 | 2.63E-03 | 2, 3 |
| **Prkcd** | protein kinase C, delta | 0.652 | 4.02E-04 | 3 |
| Fas | Fas (TNF receptor superfamily member 6) | 0.814 | 2.76E-02 | 6 |
| Ifi30 | interferon gamma inducible protein 30 | 0.836 | 4.54E-02 | 3 |
| Irf9 | interferon regulatory factor 9 | 1.108 | 1.48E-03 | 1, 3, 6, 4 |
| Irf8 | interferon regulatory factor 8 | 1.154 | 4.97E-03 | 1, 3 |
| **Socs1** | suppressor of cytokine signaling 1 | 1.162 | 1.59E-02 | 1, 3, 5, 4 |
| Fos | FBJ osteosarcoma oncogene | 1.337 | 2.17E-04 | 6 |
| **Stat1** | signal transducer and activator of transcription 1 | 1.359 | 1.51E-03 | 1, 3, 5, 6, 4 |
| Ifi35 | interferon-induced protein 35 | 1.371 | 4.93E-04 | 1, 4 |
| Ube2l6 | ubiquitin-conjugating enzyme E2L 6 | 1.404 | 2.17E-03 | 2 |
| **Irf1** | interferon regulatory factor 1 | 1.520 | 2.88E-05 | 1, 3, 6, 4 |
| **B2m** | beta-2 microglobulin | 1.586 | 2.27E-05 | 3 |
| Irf7 | interferon regulatory factor 7 | 1.732 | 6.63E-04 | 1, 2, 3 |
| **Tap1** | transporter 1, ATP-binding cassette, sub-family B (MDR/TAP) | 1.939 | 3.15E-05 | 4 |
| Isg15 | ISG15 ubiquitin-like modifier | 2.002 | 2.17E-04 | 1, 2, 4 |
| Xaf1 | XIAP associated factor 1 | 2.099 | 8.03E-09 | 1 |
| **Psmb8** | proteasome (prosome, macropain) subunit, beta type 8 (large multifunctional peptidase 7) | 2.733 | 8.58E-09 | 1, 4 |
| **Bst2** | bone marrow stromal cell antigen 2 | 3.151 | 7.37E-09 | 1 |
| H2-Q7 | histocompatibility 2, Q region locus 7 | 4.236 | 1.07E-06 | 1, 3 |
| H2-Aa | histocompatibility 2, class II antigen A, alpha | 5.489 | 4.55E-08 | 3 |

**Table 3.**
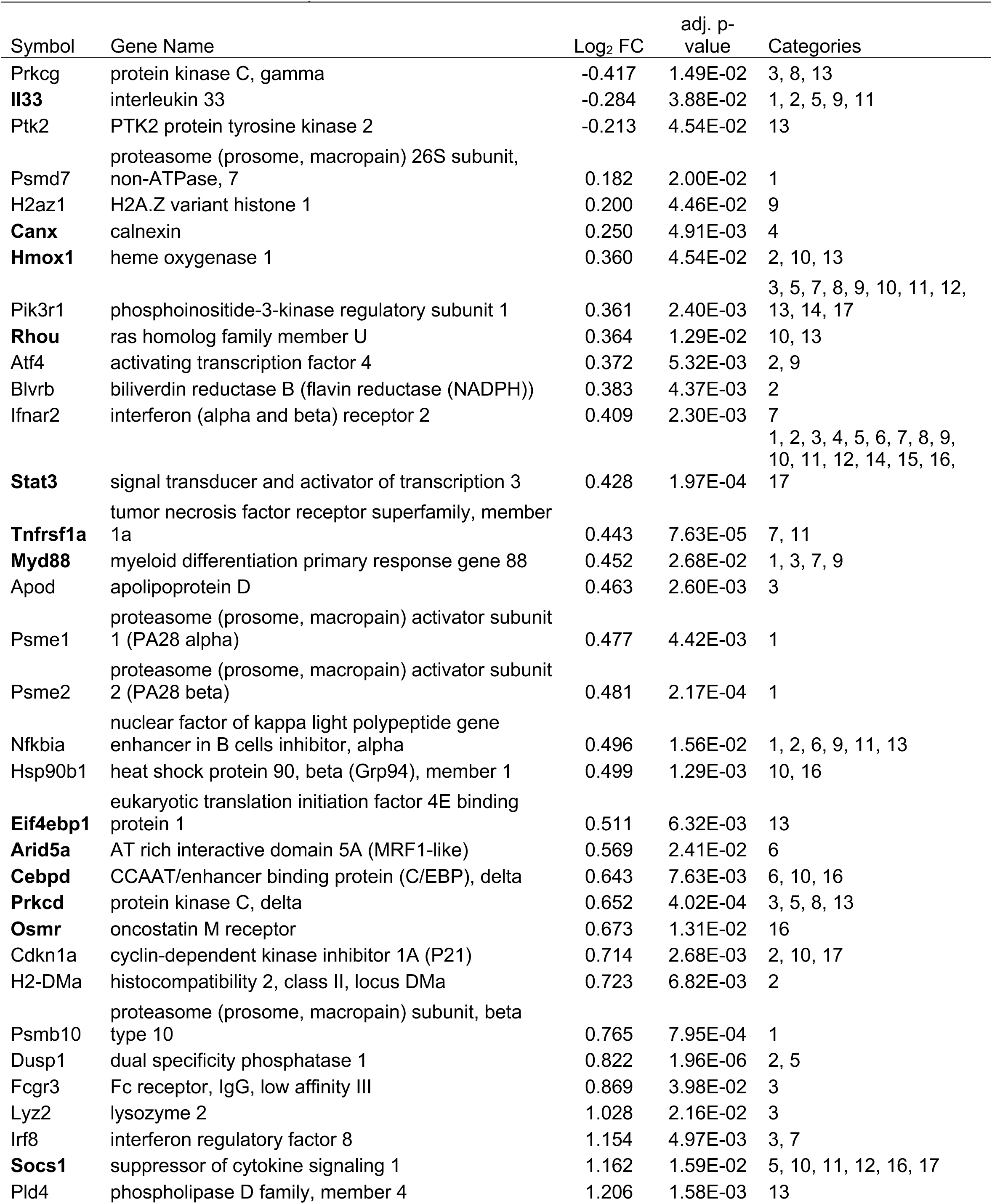
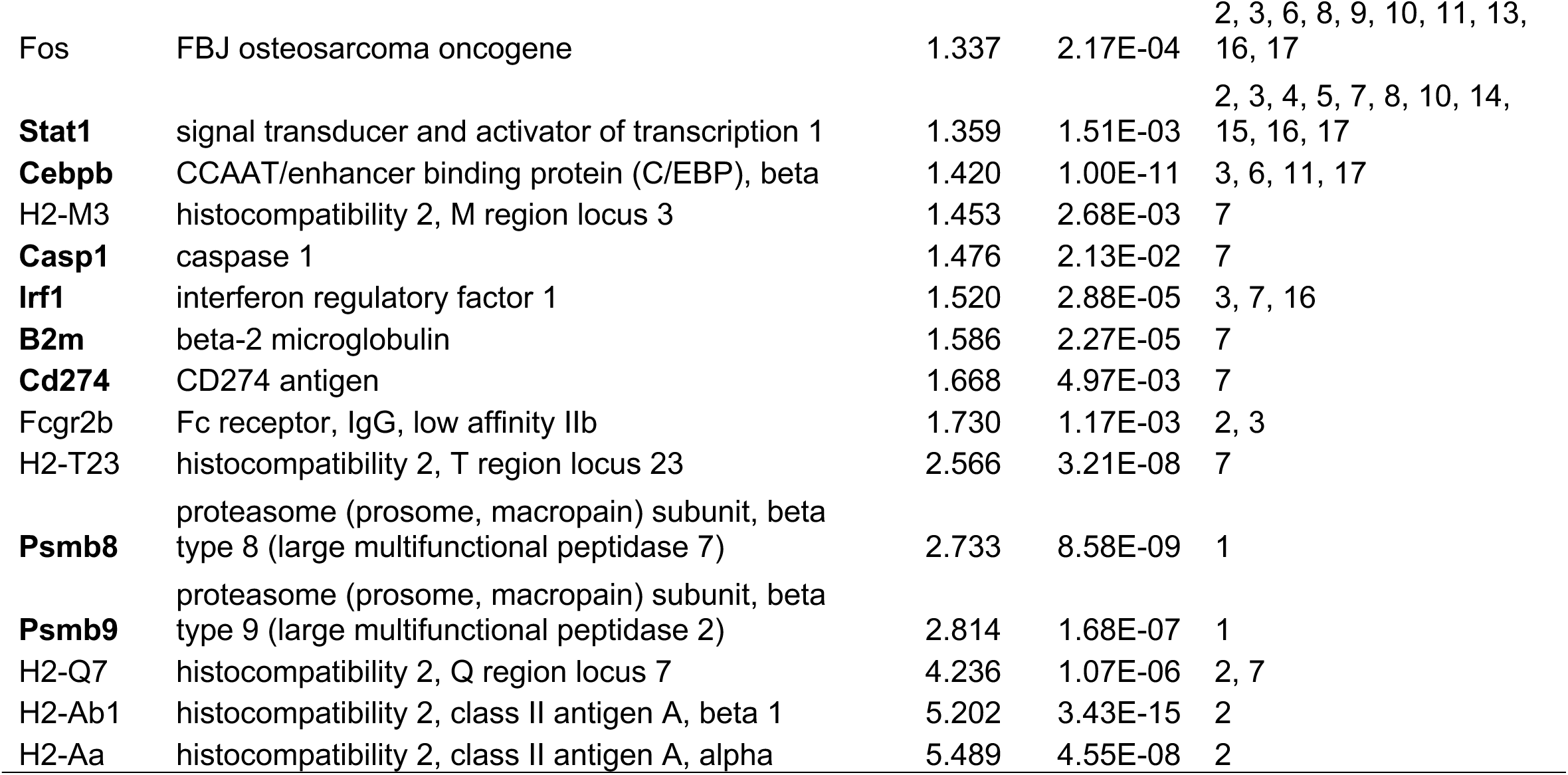
Interleukin Related Pathway Genes. Gene symbols in bold indicate the gene is enriched in the astrocyte (IP) fraction. Categories column refers to the following IPA canonical pathways: 1) Interleukin-1 family signaling; 2) IL-10 Signaling; 3) IL-12 Signaling and Production in Macrophages; 4) Interleukin-12 family signaling; 5) IL-13 Signaling Pathway; 6) IL-17A Signaling in Fibroblasts; 7) IL-27 Signaling Pathway; 8) IL-3 Signaling; 9) IL-33 Signaling Pathway: 10) Interleukin-4 and Interleukin-13 signaling; 11) IL-6 Signaling; 12) Interleukin-7 signaling; 13) IL-8 Signaling; 14) IL-9 Signaling; 15) Interleukin-9 signaling; 16) Role of JAK family kinases in IL-6-type Cytokine Signaling; 17) JAK/STAT Signaling.

Another neuroimmune function activated in Dependent animals involves interleukin signaling. “IL-10 Signaling”, “IL-27 Signaling Pathway”, and “Interleukin-4 and Interleukin-13 signaling” are among the top 30 canonical pathways in the Dependent vs non-Dependent IP comparison (Figure 3B, highlighted in yellow). There are a total of 17 significantly overrepresented canonical pathways related to interleukins (Supplemental Table 4). These pathways and significantly regulated genes are summarized in Table 3 and Figure 4B.

IPA of the significantly regulated astrocyte-enriched genes (Supplemental Table 5 and summarized in Table 1) are shown in Figure 3 C, D. Similar to what is shown in Figure 3 A,B, these categories were dominated by neuroimmune-related categories, most with positive activation z-scores. For instance, in the Dependent vs naïve comparison ‘IL-4 and IL-13 signaling’ was the most enriched pathway (Figure 3C) with a positive activation z-score and in the Dependent vs non-Dependent comparison, the second most enriched pathway is ‘IL-27 Signaling’ (Figure 3D). All IPA pathways significantly overrepresented in these comparisons are shown on Supplemental Table 4. This analysis of a subset of genes that are mostly expressed by astrocytes in the brain clearly indicates an upregulation in astrocyte neuroimmune responses further supporting the role of astrocytes in alcohol-induced neuroinflammation.

“IL-33 Signaling Pathway” was significantly overrepresented in the IPA analysis of Dependent vs non-Dependent IP samples, with a positive activation z-score indicating pathway activation. This IPA pathway contains 8 genes; 4 additional genes regulated in Dependent IP samples were identified as part of the IL-33 pathway in two recent studies (Pinto et al., 2018; Jiao et al., 2024). Of the 12 DT genes in the IL-33 signaling pathway, 4 genes (*Stat3*, *Il33*, *Myd88*, and *Tmed10*) are also enriched in astrocytes (Table 4).

**Table 4.**
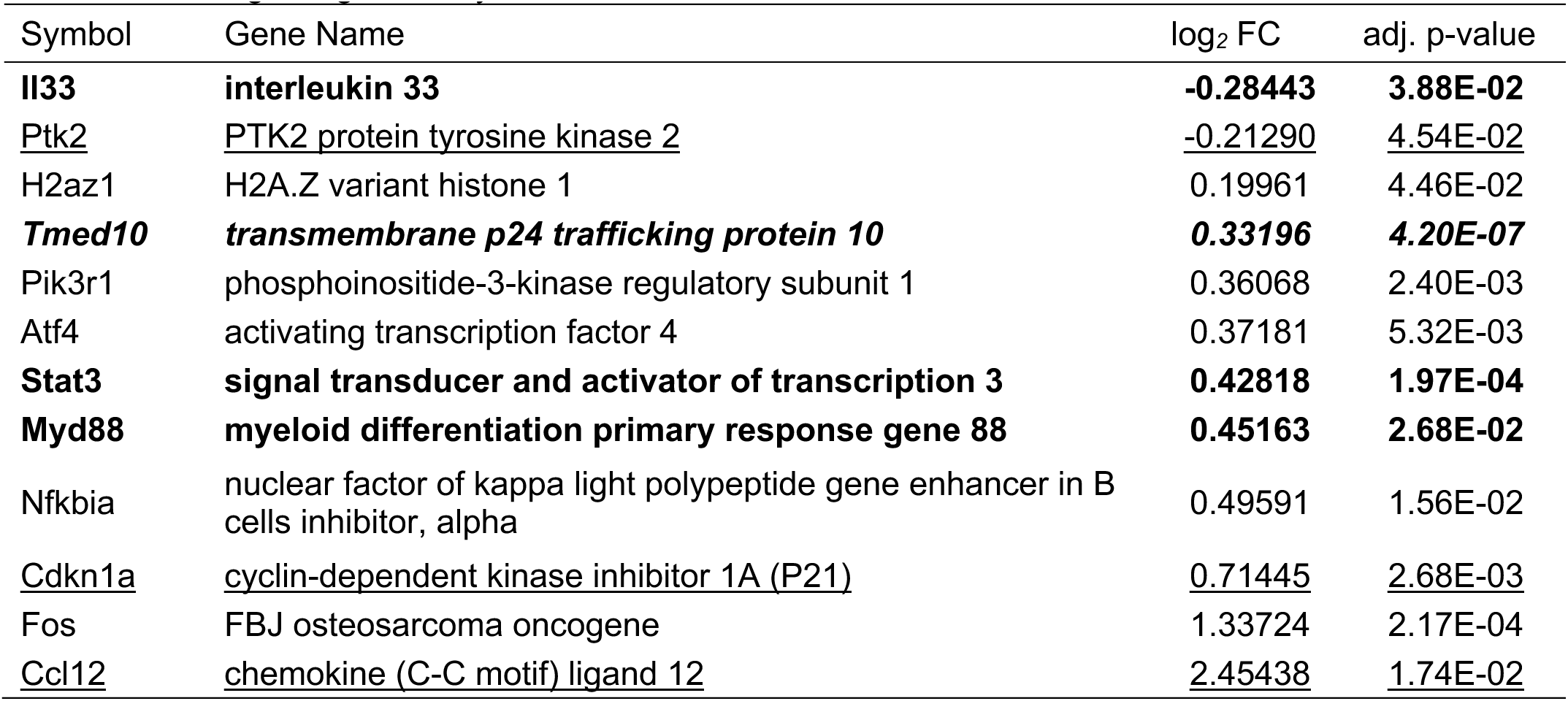
IL-33 Signaling Pathway Genes. Bold indicates gene is enriched in astrocytes. Underlined genes added based on Pinto 2018. Italics gene added based on Jiao 2024.

| Symbol | Gene Name | log<sub>2</sub> FC | adj. p-value |
| --- | --- | --- | --- |
| **Il33** | **interleukin 33** | **-0.28443** | **3.88E-02** |
| <u>Ptk2</u> | <u>PTK2 protein tyrosine kinase 2</u> | <u>-0.21290</u> | <u>4.54E-02</u> |
| H2az1 | H2A.Z variant histone 1 | 0.19961 | 4.46E-02 |
| ***Tmed10*** | ***transmembrane p24 trafficking protein 10*** | ***0.33196*** | ***4.20E-07*** |
| Pik3r1 | phosphoinositide-3-kinase regulatory subunit 1 | 0.36068 | 2.40E-03 |
| Atf4 | activating transcription factor 4 | 0.37181 | 5.32E-03 |
| **Stat3** | **signal transducer and activator of transcription 3** | **0.42818** | **1.97E-04** |
| **Myd88** | **myeloid differentiation primary response gene 88** | **0.45163** | **2.68E-02** |
| Nfkbia | nuclear factor of kappa light polypeptide gene enhancer in B cells inhibitor, alpha | 0.49591 | 1.56E-02 |
| <u>Cdkn1a</u> | <u>cyclin-dependent kinase inhibitor 1A (P21)</u> | <u>0.71445</u> | <u>2.68E-03</u> |
| Fos | FBJ osteosarcoma oncogene | 1.33724 | 2.17E-04 |
| <u>Ccl12</u> | <u>chemokine (C-C motif) ligand 12</u> | <u>2.45438</u> | <u>1.74E-02</u> |

The expression of *Il33* was significantly down-regulated in the IP fraction of Dependent mice compared to non-Dependent mice. Confirmation of down-regulation of *Il33* by RT-qPCR in Dependent vs non-Dependent TRAP NAc samples is shown in Figure 4C. RT-qPCR also confirmed that *Il33* expression is enriched in astrocytes compared to the input samples (Figure 4C).

### 3.4 EtOH dependence alters oxidative stress related pathways

A key function of astrocytes in the brain is to protect neurons from endogenous and exogenous toxins and from oxidative stress, as astrocytes are the main cell types involved in detoxification and reduction (neutralization) of reactive oxygen species primarily through the production and regulation of the glutathione system (Guitart et al., 2015; He and Hewett, 2025). Within the top 30 overrepresented canonical pathways in the Dependent vs non-Dependent IP comparison there were three pathways related to oxidative stress or glutathione that were activated: “NFE2L2 regulating anti-oxidant detoxification enzymes”, “NRF2-mediated Oxidative Stress Response”, and “Production of Nitric Oxide and Reactive Oxygen Species in Macrophages” (Figure 3B, highlighted in blue). In all IPA pathways significantly overrepresented in the Dependent vs non-Dependent IP comparison (Supplemental Table 3) there were eight pathways related to oxidative stress and glutathione; these pathways and the genes within are summarized in Table 5.

**Table 5.** Glutathione/Oxidative Stress Related Pathway Genes. Gene symbols in bold indicate the gene is enriched in the astrocyte (IP) fraction. Categories column refers to the following IPA canonical pathways: 1) Glutathione Biosynthesis; 2) Glutathione-mediated Detoxification; 3) Glutathione Redox Reactions I; 4) iNOS Signaling; 5) NFE2L2 regulating anti-oxidant/detoxification enzymes; 6) Nitric Oxide Signaling in the Cardiovascular System; 7) Production of Nitric Oxide and Reactive Oxygen Species in Macrophages; 8) NRF2-mediated Oxidative Stress Response.

| Symbol | Gene Name | Log<sub>2</sub> FC | adj. p-value | Categories |
| --- | --- | --- | --- | --- |
| **Gucy2e** | guanylate cyclase 2e | -0.754 | 4.49E-03 | 6 |
| Prkcg | protein kinase C, gamma | -0.417 | 1.49E-02 | 6, 7, 8 |
| Chrm1 | cholinergic receptor, muscarinic 1, CNS | -0.362 | 6.09E-03 | 6 |
| Cacng3 | calcium channel, voltage-dependent, gamma subunit 3 | -0.357 | 2.76E-02 | 6 |
| Ppp2r1a | protein phosphatase 2, regulatory subunit A, alpha | -0.217 | 4.59E-02 | 7 |
| Gstp1 | glutathione S-transferase, pi 1 | 0.248 | 6.69E-03 | 2, 3, 8 |
| Txnrd1 | thioredoxin reductase 1 | 0.264 | 2.16E-02 | 5, 8 |
| **Gstm1** | glutathione S-transferase, mu 1 | 0.272 | 2.84E-02 | 2, 8 |
| Gsta4 | glutathione S-transferase, alpha 4 | 0.301 | 4.54E-02 | 2 |
| Txn1 | thioredoxin 1 | 0.313 | 1.84E-02 | 5, 8 |
| Gstm6 | glutathione S-transferase, mu 6 | 0.327 | 1.80E-02 | 2, 8 |
| Dnajb11 | DnaJ heat shock protein family (Hsp40) member B11 | 0.351 | 9.11E-03 | 8 |
| **Gclm** | glutamate-cysteine ligase, modifier subunit | 0.353 | 3.92E-03 | 1, 5, 8 |
| **Ppp1r3c** | protein phosphatase 1, regulatory subunit 3C | 0.359 | 3.96E-03 | 7 |
| **Hmox1** | heme oxygenase 1 | 0.360 | 4.54E-02 | 5, 8 |
| Pik3r1 | phosphoinositide-3-kinase regulatory subunit 1 | 0.361 | 2.40E-03 | 6, 7, 8 |
| **Rhou** | ras homolog family member U | 0.364 | 1.29E-02 | 7 |
| Atf4 | activating transcription factor 4 | 0.372 | 5.32E-03 | 5, 8 |
| Gclc | glutamate-cysteine ligase, catalytic subunit | 0.378 | 1.35E-02 | 1, 5, 8 |
| **Dnajc3** | DnaJ heat shock protein family (Hsp40) member C3 | 0.393 | 8.31E-04 | 8 |
| **Dnajb9** | DnaJ heat shock protein family (Hsp40) member B9 | 0.432 | 3.86E-04 | 8 |
| **Tnfrsf1a** | tumor necrosis factor receptor superfamily, member 1a | 0.443 | 7.63E-05 | 7 |
| **Myd88** | myeloid differentiation primary response gene 88 | 0.452 | 2.68E-02 | 4 |
| Apod | apolipoprotein D | 0.463 | 2.60E-03 | 7 |
| **Ephx1** | epoxide hydrolase 1, microsomal | 0.471 | 6.68E-03 | 8 |
| **Ptges** | prostaglandin E synthase | 0.483 | 1.29E-03 | 2, 3 |
| Nfkbia | nuclear factor of kappa light polypeptide gene enhancer in B cells inhibitor, alpha | 0.496 | 1.56E-02 | 4, 7 |
| Hsp90b1 | heat shock protein 90, beta (Grp94), member 1 | 0.499 | 1.29E-03 | 6, 8 |
| **Slc7a11** | solute carrier family 7 (cationic amino acid transporter, y+ system), member 11 | 0.501 | 1.56E-02 | 5 |
| **Mgst1** | microsomal glutathione S-transferase 1 | 0.508 | 2.00E-06 | 2, 3, 8 |
| **Prkcd** | protein kinase C, delta | 0.652 | 4.02E-04 | 6, 7, 8 |
| Srxn1 | sulfiredoxin 1 homolog (S. cerevisiae) | 0.656 | 2.77E-07 | 5 |
| **Ly96** | lymphocyte antigen 96 | 0.657 | 4.76E-02 | 4 |
| Cp | ceruloplasmin | 0.845 | 6.32E-03 | 3 |
| Lyz2 | lysozyme 2 | 1.028 | 2.16E-02 | 7 |
| Irf8 | interferon regulatory factor 8 | 1.154 | 4.97E-03 | 7 |
| Fos | FBJ osteosarcoma oncogene | 1.337 | 2.17E-04 | 4, 7, 8 |
| **Stat1** | signal transducer and activator of transcription 1 | 1.359 | 1.51E-03 | 4, 7 |
| **Irf1** | interferon regulatory factor 1 | 1.520 | 2.88E-05 | 4, 7 |

### 3.5 EtOH drinking alters astrocyte homeostatic functions

In the non-Dependent vs naïve comparisons (Figure 5), the top enriched pathways include ‘Vitamin-C Transport’, ‘Plasma lipoprotein assembly, remodeling, and clearance’, and ‘Neurovascular Coupling’ for the astrocyte fraction (IP) and ‘Sleep NREM Signaling’, ‘Complement System’, and ‘CSDE1 Signaling’ (Input). In contrast with Dependent comparisons, enriched pathways in the comparison of non-Dependent with ethanol-naïve mice did not include neuroimmune terms. Importantly, while ethanol vapor exposure induced activation of several neuroimmune pathways involving interferon and cytokine signaling and oxidative stress in astrocytes suggesting a transition toward astrocyte reactivity (Figure 3), the changes induced by alcohol drinking are more in line with the physiologically homeostatic role of astrocytes. In support of this observation, the activation z-score of enriched pathways in both the astrocyte fraction and the bulk-tissue fraction shows little activation or deactivation suggesting homeostatic regulation, i.e. compensatory changes to pathway perturbations (Figure 5).

**Figure 5.**
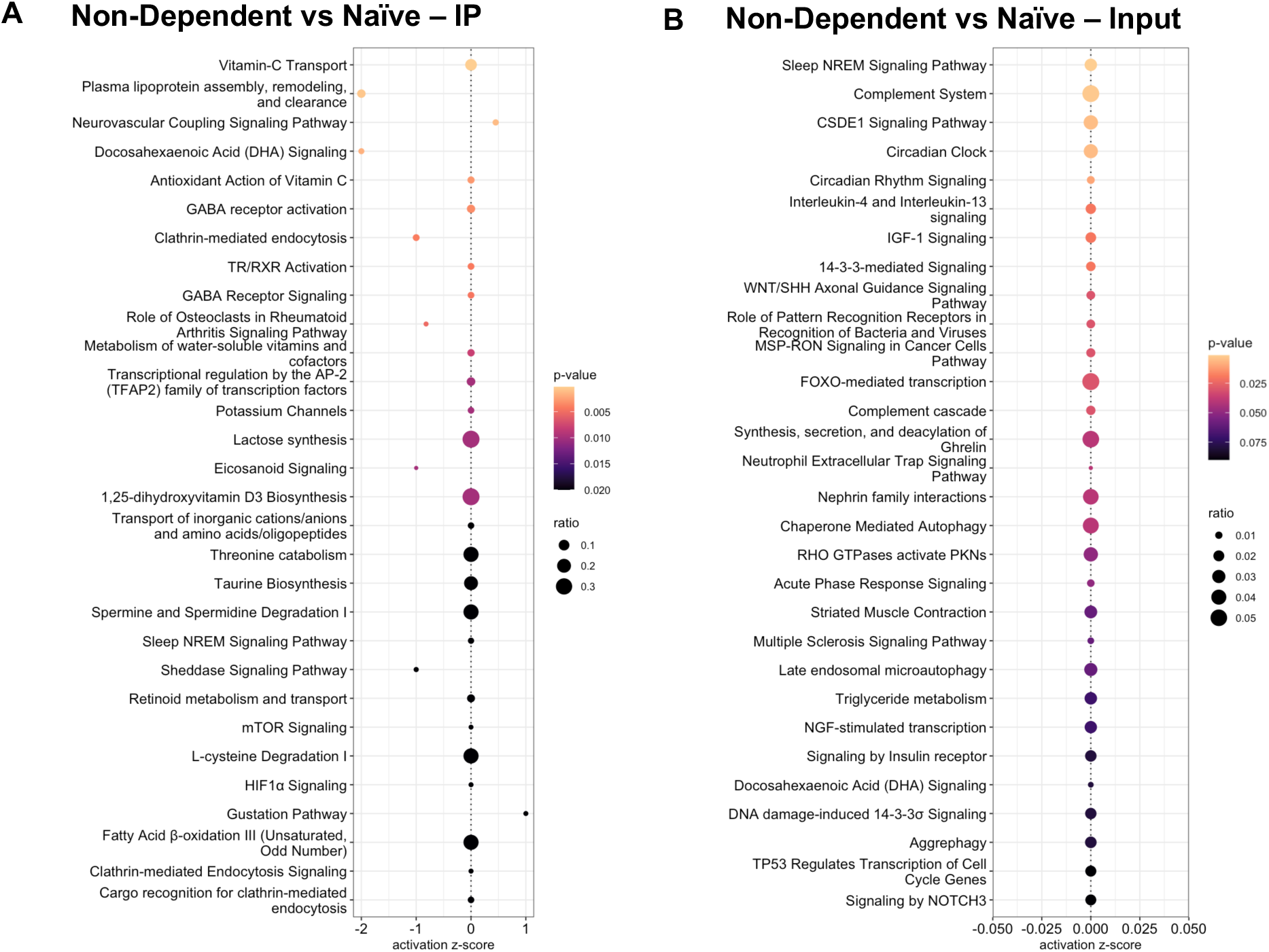
Top 30 enriched canonical pathways from IPA of non-Dependent vs naïve IP (**A**) and Input (**B**) comparisons.

## Discussion

The objective of this study was to understand how chronic alcohol and EtOH dependence in particular alters NAc astrocytes. The main findings of this study are: EtOH dependence causes a strong neuroimmune response in astrocytes including activation of interferon and interleukin pathways while EtOH drinking alone caused homeostatic responses in NAc astrocytes but did not show evidence of a neuroimmune activation.

### EtOH dependence induced astrocyte reactivity and neuroinflammatory signaling

Astrocytes have been implicated in several brain pathologies and are targets for potential therapeutics (Almad and Maragakis, 2018; Gorshkov et al., 2018; Valori et al., 2019). Researchers have started exploring the roles of astrocytes in drug and alcohol use disorders, though, in the case of AUD, considerable research still needs to be done, as highlighted in a recent review (Erickson et al., 2018; Linker et al., 2019; Skupio et al., 2020; Guizzetti et al., 2025). Astrocytes in the healthy brain interact with all cell types and play many roles required for normal brain function but, in response to CNS insults, can become reactive. Reactive astrocytes provide adaptive responses to initial brain insults and the attenuation of astrocyte reactivity can exacerbate tissue injury and worsen diseases’ outcomes (Bush et al., 1999; Faulkner et al., 2004; Kraft et al., 2013; Liu et al., 2014). Reactive astrocytes, however, can also exert maladaptive effects through inappropriate gain of function that may lead to excessive inflammation. For instance, chronic astrocyte exposure to reactivity triggers during autoimmune or neurodegenerative disorders leads to increased inflammatory responses (Sofroniew, 2015; Wheeler et al., 2020).

To determine if the NAc astrocytes exposed to EtOH become reactive, i.e., pro-inflammatory, in the CIE-2BC exposure model we compared DT genes from astrocytes to known markers of reactive astrocytes (Escartin et al., 2021). In agreement with the idea that EtOH causes astrocyte activation we see up-regulation of *Stat3* and *Tspo* in the Dependent vs non-Dependent IP comparison; in addition, *Vim*, *Gfap*, and *Mt1* were up-regulated in the Dependent vs naïve IP comparison (Supplemental Table 6). All 5 of these genes were highlighted as markers for reactive astrocytes with increased expression an indicator of astrocyte reactivity (Escartin et al., 2021) confirming the relationship of EtOH dependence causing astrocyte reactivity.

Interferon alpha/beta and gamma pathways, and specific interferon-regulatory genes such as *Irf1* and *Irf7*, were up-regulated in Dependent mice compared to non-Dependent and naïve mice. This observation extends prior work demonstrating that EtOH dependence upregulates interferon signaling in other brain regions and species. Type 1 interferon genes (IFN-alpha/beta) were upregulated in the prefrontal cortex (PFC) (Farris et al., 2020) and nucleus of the solitary tract (Grantham et al., 2024) of mice made dependent in the 2BC+CIE model, and in the PFC of mice made dependent by CIE vapor alone (Erickson et al., 2019). Moreover, interferon-related gene expression was also elevated in the PFC and the NAc of human subjects with alcohol dependence (Kapoor et al., 2019; Drake et al., 2020). Conversely, systemic activation of interferon signaling leads to increased voluntary EtOH intake (Grantham et al., 2020; Lovelock et al., 2022; Allard et al., 2024), suggesting that the upregulation of interferon pathways during EtOH dependence plays a causal role in the escalation of EtOH intake.

While many prior studies have led to the identification of interferon signaling as a conserved cross-species mechanism underlying alcohol use disorder (Friske et al., 2025), interleukin signaling pathways are less commonly identified. Our results point to the specific regulation of interleukin pathways in astrocytes (in the IP fraction), with fewer significant changes noted when examining the input. This may explain why prior studies that did not examine cell-type specific differences may have not observed such prominent effects on interleukin signaling pathways. Although interleukin signaling pathways showed positive activation z-scores, we observed a dependence-related decrease in *Il33* in particular. The down-regulation of *Il33* may represent a compensatory reaction of astrocytes to offset the overactivation of pro-inflammatory pathways. Interestingly, *Il33* was downregulated and *Myd88* was upregulated in Dependent animals only in the NAc, while they were not altered in the amygdala and in the medial PFC (data not shown), indicating that altered *Il33* signaling specifically in the NAc may play an important role in the development of alcohol dependence.

Intriguingly, IL-33 signaling in the brain regulates glutamatergic activity and promotes excitatory synapse plasticity (Nguyen et al., 2020; Wang et al., 2021). This suggests the possibility that changes in astrocyte production of IL-33 and altered activity of IL-33 may contribute to the glutamatergic plasticity induced by ethanol dependence. Independently of any role for IL-33, we also observed other evidence that astrocytes may participate in ethanol dependence-elicited glutamatergic and synaptic adaptations. “Glutaminergic Receptor Signaling Pathway (Enhanced)” was overrepresented in the IPA canonical pathway analysis comparing Dependent versus non-Dependent DT genes (p-value = 0.00428), with a positive activation z-score of 1.51 (Supplemental Table 4). Astrocytes are also involved in the complement system of the brain and respond to and provide key signals in this pathway (Schartz and Tenner, 2020). “Complement System” was significantly overrepresented in the Dependent vs non-Dependent IP fraction comparison (p-value = 1.6×10^−4^) with a positive activation z-score and included the genes *Serping1*, *C1qa*, *C1qb*, *C1qc*, and *C4b* (Supplemental Table 4). Activation of the complement system is of interest as it modulates the immune response, including neuroinflammation, but is also involved in synapse pruning (Gomez-Arboledas et al., 2021).

An important function of astrocytes in the brain is the maintenance of proper redox balance through the production and regulation of the glutathione system, and of antioxidant proteins including metallothionein (Walz and Mukerji, 1988; Chen et al., 2001; Escartin et al., 2021). Reduction of reactive oxygen species (ROS) and reactive nitrogen species (RNS) by astrocytes is essential as neurons have limited ability to protect themselves from oxidative stress while at the same time requiring high levels of oxidative metabolism for energy, and astrocyte dysfunction leading to increased oxidative stress in the brain can lead to neuroinflammation (Chen et al., 2020). In Dependent mice, we see significant enrichment in several pathways relating to glutathione and ROS/RNS in NAc astrocytes (Figure 3, Table 5), suggesting that astrocytes are compensating for increased oxidative stress due to ethanol itself or to the effect of ethanol on neurons and other brain cells with upregulation of the glutathione system, though overactivation of this system may lead to antioxidant depletion and oxidative stress, as indicated by the activation of pathways involved in nitric oxide and reactive oxygen species production (Figure 3,Table 5).

### Homeostatic astrocyte adaptations to EtOH drinking

A benefit of the CIE-2BC model of EtOH dependence is the ability to compare different qualitative levels of EtOH experience. The non-Dependent vs naïve comparisons show the effects of voluntary moderate EtOH drinking while the Dependent vs non-Dependent comparison shows the effects of excessive EtOH exposure resulting in escalating drinking, a model of dependence. Consistent with this, we found no significant neuroimmune-related categories in the non-Dependent vs naïve IPA (Figure 5). Of note, regulation of specific inflammatory pathways in CIE-2BC mice compared to naïve and non-Dependent mice is likely due to a combination of the greater EtOH intake and vapor exposure in these mice, with escalated EtOH intake characterizing the dependence phenotype. For example, interferon pathway genes *Irf1*, *Irf7*, and *Irf9* and interleukin pathway genes *Casp1* and *Il33* showed differential expression and/or translation when comparing CIE-2BC mice to naïve or non-Dependent mice, but not when comparing non-Dependent to naïve mice. The concurrence in the identification of interferon signaling between our study and those using human postmortem brain tissue (which compare individuals with alcohol use disorder to controls that drink some alcohol) demonstrates the utility of the mouse CIE-2BC model and comparison with 2BC voluntary drinking for modeling human alcohol use disorder and identifying dependence-specific changes in gene expression.

Voluntary EtOH drinking, on the other hand, induces changes that we interpreted as homeostatic, based on the observations that fewer genes are regulated in the comparison between naïve vs EtOH-drinking (non-Dependent) animals than in the comparison between Dependent vs. non-Dependent animals, and that there are very few overlapping genes in the two sets of comparisons. Even more important, EtOH drinking in non-Dependent mice does not induce neuroimmune activation and the pathways that are significantly overrepresented in IPA analysis have z-scores around 0, indicating that EtOH drinking does not induce activation or inhibition of specific pathways, but rather that changes in gene translation induced by ethanol drinking are compensated by homeostatic changes in other genes of the same pathway resulting in a lack of overall perturbation of these pathways.

## Conclusions

Our results indicate that astrocytes are major players in the neuroimmune response elicited by the CIE-2BC model of EtOH dependence. While neuroimmune activation was reported by us and others after exposing animals to EtOH vapor using the CIE-2BC model (Warden et al., 2020; Patel et al., 2021; Varodayan et al., 2023), the contribution of astrocytes to neuroimmune responses has never been characterized before. We show that there is an activation of immune function-related pathways in astrocytes, particularly interferon and interleukin signaling pathways.

## Acknowledgements

Support for this study was provided by National Institute on Alcohol Abuse and Alcoholism grants: U01AA013498, P60 AA006420, R37 AA017447, R01AA021491, R01AA029841, and the Schimmel Family Chair (MR). U01AA016651, R01AA030256, the Billi Terresa Cowden Endowment administered by the Waggoner Center for Alcohol and Addiction Research at The University of Texas at Austin (R.A.M.).

This study was supported by NIH U01AA029965, P60AA010760, R01AA029486, and R21DA060442, and VA Merit Review Award I01BX001819, and by facilities and resources at the Portland VA Health Care System to M.G. The contents of this article do not represent the views of the United States Department of Veterans Affairs or the United States government.

## Notes

### Competing Interest Statement

The authors have declared no competing interest.

### Summary of Updates

Figure 4 was repeated and Figure 5 was omitted in the original submission.

